# Demographic compensation buffers population decline under variable grazing intensities in alpine grasslands

**DOI:** 10.64898/2026.07.18.739300

**Authors:** Hai-Tao Miao, Roberto Salguero-Gómez, Yukun Hu, Xiaofang Wang, Zhenhua Zhang, Jin-Sheng He, Shou-Li Li

## Abstract

Identifying optimal grazing intensities for sustainable population growth is crucial for informing management strategies. Community-level studies frequently find that biodiversity peaks at intermediate grazing intensities, known as the ‘Intermediate Disturbance Hypothesis’. However, whether this hypothesis applies to population-level performance remains untested. Our stochastic integral projection models, parameterized with five-year demographic data of two co-occurring species, *Morina chinensis* and *Deyeuxia flavens*, on the Tibetan Plateau grasslands., indeed show a hump-shaped response in stochastic population growth rate (λ_S_) to grazing intensities, providing empiral support for the ‘Intermediate Disturbance Hypothesis’ at population level. Furthermore, populations with demographic compensation among vital rates are better able to buffer temporal variation in annual population growth rate and exhibit a much smaller decline in λ_S_ under heavy grazing. Our study provides mechanistic insights into demographic processes driving population dynamics across grazing levels, thereby better informing grazing management strategies.

## INTRODUCTION

Livestock grazing is a major anthropogenic disturbance to the global biodiversity (Filazzola et al. 2020; Gao & Carmel 2020). The effects of grazing on species diversity largely depend on grazing intensity (Eldridge et al. 2018; Filazzola et al. 2020). Many studies at community level have shown that grazing generally increases species diversity at low or intermediate intensity, but decreases it at high intensity, which is well-known as ‘Intermediate Disturbance Hypothesis’ (IDH) (Connell 1978; Huston 1979; Hixon & Brostoff 1983). Empirical and modeling evidence suggests that species diversity in response to grazing intensity relies on the comprehensive grazing effects on survival, growth, and fecundity over life history cycle (Huston 1979; Hacker & Gaines 1997; Hart & Jenkins 2023), and ultimately the intrinsic population growth rates of species in communities (Huston 1979; Fox 2013; Huston 2014; Hart & Jenkins 2023). Therefore, a thorough understanding of plant diversity-grazing intensity relationship requires downscaling the grazing effects at the community level to the demographic process at population scale (Huston 1979). However, it remains unknown whether IDH still holds for the relationship between the intensity of grazing and the population performance.

Understanding how population viability respond to grazing requires demographic studies across experimentally controlled gradients of grazing intensity. Currently, most demographic studies only examined population viability under the business as usual grazing conditions *vs*. grazing exclusion conditions, without an explicit consideration of grazing intensities (Martorell 2007; Li et al. 2013; Aschero et al. 2016; Larios & Hallett 2022). Furthermore, although some demographic studies have examined the effects of grazing intensity on population viability, key contextual factors—such as the habitat quality (Knight et al. 2009; Evju et al. 2010, 2011), grazing history (Hegland et al. 2010) and grazing season (Bastrenta et al. 1995; Johansen et al. 2016)—are often uncontrolled. These factors frequently vary among treatments within the same study, making it difficult to disentangle the effects of grazing intensity on population dynamics. Thus, to thoroughly understand the effects of grazing on population viability, it is essential to conduct a manipulative experiment with well-controlled grazing contexts across a gradient of grazing intensities.

Whether a species can maintain viable populations under different grazing intensities depends on the synthesized responses of all the demographic rates (*e*.*g*., survival, growth, and fecundity). The response of vital rates to grazing intensity may vary in magnitude, and sometimes in opposite directions (Evju et al. 2010; Hegland et al. 2010). If the decline in some demographic rates at a given grazing intensity is offset by an increase in others, the overall population-level effects at that grazing intensity could be alleviated or cancelled out, a phenomenon known as ‘demographic compensation’ (Villellas et al. 2015; Sheth & Angert 2018). Consequently, the presence of demographic compensation may decrease variation in population growth across a variety of grazing levels and even prevent population collapse under heavy grazing conditions. For example, Evju *et al*. (2010) found that the alpine plant *Viola biflora* L. was able to compensate for grazing-induced reductions in fecundity through increased growth. Such compensation among vital rates stabilized population growth of *V. biflora* across the grazing intensity and forestalled its population crash at high grazing intensity (Evju et al. 2010). Conversely, populations lacking demographic compensation are more prone to decline with the increasing grazing intensity (Hegland et al. 2010). Thus, assessing demographic compensation across grazing intensities may provide deeper insights into population dynamics and better inform sustainable grazing management.

Long-term population viability under grazing also depends on the mean and temporal variance of the annual population growth rate in a stochastic environment (Jacquemyn et al. 2012; Larios & Hallett 2022). Life-history theory consistently predicts that, given the mean annual population growth rates remain constant, increasing temporal variation may lead to a reduction in the long-term stochastic population growth rate (Tuljapurkar 1982; Pfister 1998; Hilde et al. 2020). Therefore, predicting the long-term viability of grazed populations under environmental change requires integrating both the mean and temporal variation of vital rates within a stochastic framework (Jacquemyn et al. 2012; Larios & Hallett 2022). While previous empirical studies have extensively examined the impact of grazing-induced changes in mean vital rates on population maintenance (Bastrenta et al. 1995; Evju et al. 2010; Li et al. 2013; Larios & Hallett 2022), the role of temporal variation has received far less attention. This gap is largely due to the limited availability of long-term demographic data from grazed populations (but see refs. Farrington et al. 2009; Evju et al. 2010; Larios & Hallett 2022). The failure to disentangle the respective effects of the mean and temporal variation in vital rates under grazing largely limits our understanding of the grazed population dynamics in variable environments.

Alpine grasslands are among the most vulnerable ecosystems worldwide to anthropogenic disturbances (Yao et al. 2012; Wang et al. 2022). Intense overgrazing has led to widespread grassland degradation on the Tibetan Plateau (Bardgett et al. 2021; Wang et al. 2022). A key ecological mechanism underlying grassland degradation is selective foraging by herbivores, which drives shifts in plant community composition and structure (Cingolani et al. 2005; Díaz et al. 2007; Oñatibia et al. 2023). Increasing evidence suggests that herbivore selectivity—largely shaped by interspecific variation in plant palatability—acts as a trait-based ecological filter that governs community assembly (Cingolani et al. 2005; Díaz et al. 2007; Oñatibia et al. 2023). Plant species differing in palatability may exhibit contrasting demographic responses to grazing. Many palatable species appear to tolerate herbivory through compensatory regrowth responses such as accelerated vegetative growth and enhanced resource allocation to reproduction (Belsky 1986; Augustine & McNaughton 1998). In contrast, unpalatable species exhibit more limited compensatory responses to grazing (Augustine & McNaughton 1998; Teague et al. 2016). However, the demographic mechanisms linking plant palatability to population dynamics remain poorly understood. Clarifying how plant palatability shapes vital rates is critical for predicting long-term population dynamics under varying grazing intensity. Moreover, incorporating such demographic information into population models can help disentangle the mechanisms by which herbivore selectivity drives vegetation change and grassland degradation.

Here, we examine the demographic responses of two alpine herbaceous species with different palatability, *Morina chinensis* (Bat.) Diels (less palatable species) and *Deyeuxia flavens* Keng. (more palatable species), to different levels of grazing intensity on an alpine grassland of the Tibetan Plateau. We conducted annual censuses over a five-year period from 2019 to 2023. We utilized the demographic data to parameterize stochastic integral projection models (stochastic IPMs), which allow for incorporation of temporal fluctuations in vital rates under stochastic environments (Rees & Ellner 2009; Ellner et al. 2016). Additionally, we employed stochastic life table response experiment (SLTRE) analysis to assess the contributions of different demographic rates to the observed differences in stochastic population growth rates under different grazing intensities (Davison et al. 2010; Villellas et al. 2013). Specifically, we used the stochastic population models to test the following hypothesis: i) The hump-shaped response of alpine plants to grazing intensity can be tested using estimates of stochastic population growth rates, as predicted by the IDH; ii) *D. flavens* is expected to exhibit demographic compensation among vital rates across grazing intensities, thereby buffering population growth. In contrast, *M. chinensis* is not expected to exhibit such compensation and is therefore more vulnerable to population decline under higher grazing pressures; iii) In a stochastic environment, the effects of changes in mean vital rates on grazing-induced shifts in population growth are comparable to the effects of changes in temporal variances of vital rates. This finding will contribute to our understanding of how livestock grazing influence long-term population growth and the potential mechanisms that may allow species to adapt to different grazing conditions in fluctuating environments.

## MATERIALS AND METHODS

### 2.1 Study sites and experimental design

The study was conducted at the Haibei National Alpine Grassland Ecosystem Research Station (37°36′ N, 101°19′ E, 3,125 m a.s.l), located in the northeastern of the Tibetan Plateau, Qinghai Province, China (Figure 1a). The climate is characterized by short, cool summers and long, severely cold winters. The mean annual temperature is -1.1 **□**, with the highest monthly temperature of 10.4 **□** occurring in July and the lowest monthly temperature of -14.6 **□** occurring in January. The mean annual precipitation is 481 mm, more than 80% occurring as rain during growing season, from May to September. The study area is covered by alpine meadow (Figure 1a), and the vegetation in our study site is dominated by perennial herbaceous plant, such as *Stipa aliena, Poa crymophila*, and *Kobresia humilis*. Livestock husbandry is the major land use due to the extensive area of rangeland, with livestock number nearly doubling since 1978 (Chen et al. 2013). However, improper grazing management, particularly overgrazing resulting from rangeland privatization, is believed to lead to severe rangeland degradation or even desertification (Wang et al. 2022).

**Figure 1.**
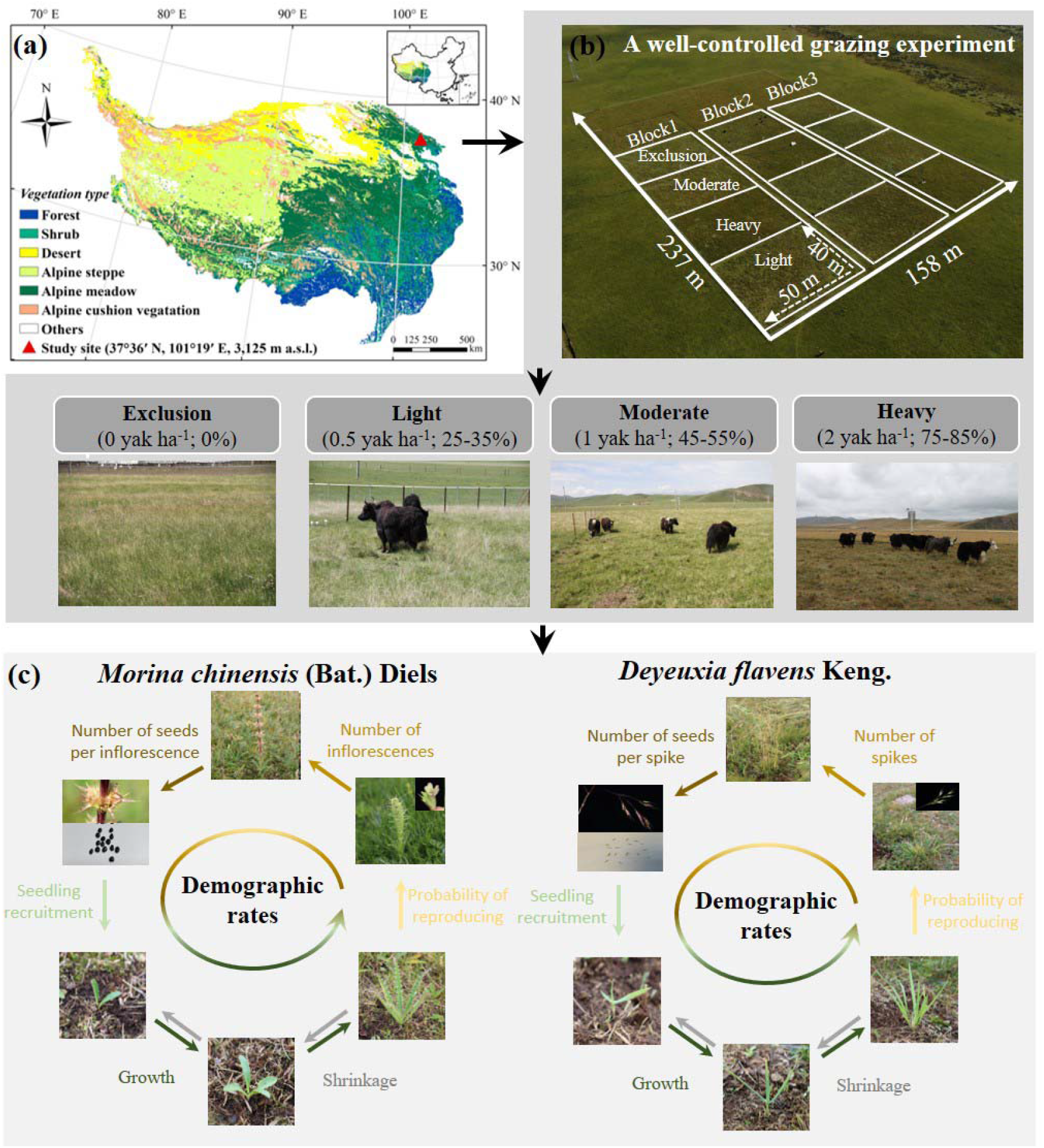
Two co-occurring species, *Morina chinensis* (Bat.) Diels and *Deyeuxia flavens* Keng. under a well-controlled grazing experiment on the Tibetan Plateau. (a) The geographical location of the study site on the Tibetan Plateau. (b) The well-controlled grazing experiment site. We established a grazing manipulative experiment with an area of 158 m × 237 m in 2019, using a randomized block design. Each block comprised four grazing intensity: grazing exclusion (Exclusion), light grazing (Light), moderate grazing (Moderate), and heavy grazing (Heavy). Each grazing intensity had three plots with an area of 40 m × 50 m. The grazing intensity was determined using the rangeland carrying capacity calculation method published in the agricultural industry standard NY/T 635–2015 (Ministry of Agriculture, People’s Republic of China). (c) The demographic processes for the two study species, *M. chinensis* and *D. flavens*, at the study site.

A large grazing manipulative experiment was established in 2019 within an area of 158 m × 237 m (Figure 1b; Wang et al. 2023). A randomized block design was employed in the experiment with four grazing intensities and three blocks (Figure 1b). Each grazing intensity included three plots with an area of 40 m × 50 m, and one/two permanent 1 m × 4 m subplots within each plot was randomly established for further demographic censuses (Figure 1b). The grazing intensity was determined using the rangeland carrying capacity calculation method published in the agricultural industry standard NY/T 635–2015 (Ministry of Agriculture, People’s Republic of China). The grazing intensity includes: (i) grazing exclusion (0 yak ha^-1^: feeding rate 0%, Exclusion), (ii) light grazing (0.5 yak ha^-1^: feeding rate 25-35%, Light), (iii) moderate grazing (1.0 yak ha^-1^: feeding rate 45-55%, Moderate), (iv) heavy grazing (2.0 yak ha^-1^: feeding rate 75-85%, Heavy). Rotational grazing was conducted annually from June to September, with each block grazed by yaks in rotation for approximately 4 days.

### 2.2 Study species and demographic study

To compare the effects of grazing intensity on the population performance of plant species with different palatability inhabiting the same habitats, we selected two co-occurring herbaceous species, *Morina chinensis* and *Deyeuxia flavens*. Both species are common species in the study area, but differ in palatability: *M. chinensis* is a less palatable, whereas *D. flavens* is a more palatable to yak (Appendix S1: Figure S1). Moreover, *M. chinensis* is a perennial forb, with linear-lanceolate leaf blade and 4-9 dense verticillasters in an erect growth stem at the reproductive stage. Conversely, *D. flavens* is a perennial herb, with erect culm and flat leaf blade and 1-20 loose panicles at the reproductive stage. Both species have similar life history: they sprout new leaves annually, regrow in April-May, grow flower stems from leaves in June-July, and set fruit in August-September, with the aboveground part of the plants withering in winter (Figure 1c). Seed recruitment generally occurs through spring germination in the following year for both species. The onset of sexual reproduction typically occurs in the fourth year for *M. chinensis* and the eighth year for *D. flavens* (as estimated by IPMs; Table 1). The plants of *D. flavens* have a high capacity for asexual reproduction through basal tillering, with new ramets typically forming in close proximity to the genet and initiating vegetative reproduction in the following years (as observed during our study periods; Table 1). Both species are widespread on the grasslands of the Tibetan Plateau, where *D. flavens* is an important forage plant and *M. chinensis* is an important medicinal plant.

**Table 1.** The summary of key traits (palatability, reproductive strategy, mean age at first reproduction, mean lifespan from reproduction, and lifespan) for the two co-occurring alpine herbaceous species, *Morina chinensis* and *Deyeuxia flavens*, on the Tibetan Plateau.

| **Traits** | ***Morina chinensis*** | ***Deyeuxia flavens*** |
| --- | --- | --- |
| Palatability | Less | More |
| Reproductive strategy | Sexual | Sexual and vegetative |
| Mean age at first reproduction (years) | 4.59 [3.92, 5.25] | 8.32 [7.09, 9.82] |
| Mean lifespan from reproduction (years) | 5.11 [4.51, 5.74] | 8.84 [7.66, 10.28] |
| Lifespan (years) | 10.21 [8.00, 12.00] | 21.67 [15.00, 28.00] |

Note: values in brackets refer to 95% confidence intervals (CI).

To examine the effect of grazing intensity on demographic rates, annual censuses were conducted in September for both species from 2019 to 2023. In the first census, we measured plant height, the length of the longest leaf and the total number of leaves, which are commonly used as proxies for individual-level state variables (Ellner et al. 2016; Struckman et al. 2019). By comparing the variance explained (*R*^*2*^) values of the three-variable regression models for each demographic rate (Nakagawa & Schielzeth 2013), the natural logarithm of current plant height was the most strongly related to individual vital rates in both species and was used as a measure of plant size. We also recorded the reproductive status of each individual for both species in each subplot, and counted the number of spikes or inflorescences produced by each reproductive individual. To estimate the seed production, we randomly selected 96 spikes for *D. flavens* and 155 inflorescences for *M. chinensis* close to our experimental site in 2019, and then brought them back to the laboratory to count the number of seeds. Upon the first measurement, we tagged each individual with a stainless steel label and recorded its Cartesian coordinates within the subplots to allow tracing through time. In the subsequent annual census from 2020 to 2023, we checked the survival status of all tagged individuals and remeasured size and the aforementioned reproductive characteristics on surviving ones. At that point, we also located and measured new recruits within our subplots. To avoid misidentifying individuals as dead due to complete removal of above-ground tissues by yaks, we rechecked all previously recorded dead individuals during each annual census from 2021 to 2023. The sampling effort resulted in a total of 1509 individuals for *D. flavens* and 2294 individuals for *M. chinensis*.

### 2.3 Environmental variation

Substantial interannual variation in demographic rates was observed across the study years, motivating the incorporation of environmental variability into population modeling. For instance, the proportion of reproductive individuals in grazed populations of *M. chinensis* declined by 95% during 2021–2022, while the proportion of dead individuals in grazed populations of *D. flavens* doubled during 2022–2023 (Appendix S1: Figure S2-S3). To evaluate how environmental fluctuations affect population dynamics under different grazing intensities, each census period (*i*.*e*., environmental condition) were classified into one of three environmental states: dry, normal, and wet years. This classification was based on total annual precipitation, a key proxy for plant population dynamics (van de Pol et al. 2016; Wang et al. 2020). Specifically, dry years had total precipitation lower than 1 standard deviation, while wet years had total precipitation greater than 1 standard deviation (Xu et al. 2020). To accurately capture the demographic responses to environmental variation, total annual precipitation was calculated by summing the precipitation from October at year *t* to September at year *t + 1* (van de Pol et al. 2016; Wang et al. 2020). Based on the long-term precipitation records spanning 1980-2023, normal years were observed from 2019 to 2021, followed by a wet year in 2021-2022 and a dry year in 2022-2023 (Appendix S1: Figure S4). To incorporate temporal autocorrelation among environmental conditions into stochastic simulations of population dynamics, we defined a discrete Markov chain consisting of three environmental states (Appendix S2). Transition probabilities between states were estimated from long-term precipitation records (Appendix S1: Figure S5).

### 2.4 Vital rates estimation

To evaluate how grazing intensity and initial plant size affect demographic rates, we employed mixed-effects models relating each demographic rate to the natural logarithm of current plant height and grazing intensity. We collected demographic data from five consecutive annual censuses across different grazing intensities, and then pooled the data across years and grazing treatments. These data were used to fit models for six key demographic processes: (a) probability of survival, (b) size change (*e*.*g*., growth and shrinkage), (c) probability of reproduction, (d) the number of inflorescences or spikes produced per reproductive plant, (e) seedling recruitment, and (f) recruit size. Each demographic process was modeled using an appropriate statistical distribution that reflected the biological nature of the response (Appendix S2).

To identify the best predictors of grazing intensity and initial plant size for each demographic rate, we constructed multiple mixed-effects models (Appendix S1: Table S1). Grazing intensity, the natural logarithm of current plant height, and their interaction were treated as fixed effects, with year as a random effect to account for temporal environmental variability (Ellner et al. 2016; Tredennick et al. 2018). Because some plots lacked individuals of particular life stages (*e*.*g*., seedlings, dead, or reproductive plants), we aggregated data across plots to avoid spurious random plot effects (Bolker et al. 2009). To select the best-supported model for each vital rate, model selection was performed via Akaike information criterion, with convergence and overfitting diagnostics applied to ensure robustness (Appendix S1: Table S2; Burnham & Anderson 2004; Barr et al. 2013; Matuschek et al. 2017). The fixed effects in the best-supported vital rate models were tested using the Wald chi-squared test, and random year effects were assessed using the Likelihood ratio test (Appendix S1: Table S3; Yang 1998; Fox & Weisberg 2019).

### 2.5 Stochastic population model

To quantify the effects of grazing intensity on long-term population dynamics of plant species, we constructed stochastic Integral Population Model (IPMs) of both study species (Rees & Ellner 2009). IPMs use information on how an individual’s state influences vital rates to project population change in discrete time (Easterling et al. 2000). Each IPM was parameterized with multi-year estimates of vital rates derived from the best-supported vital rate models. In our IPMs, the continuous state variable was the natural logarithm of current plant height log_*e*_(height) and the discrete time step (from *t* to *t* + 1) corresponded to one year. The time-varying size-structed IPM is

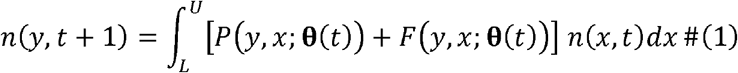

where *L* and *U* are the lower and upper bounds on the range of possible heights log_*e*_(height) for all grazing intensities, and **θ**(*t*) is a vector of year-specific parameters, assumed to vary over time. The variable *n*(*x,t*) is the distribution of log_*e*_(height) *x* at time *t*, and *n*(*y,t* + 1) is the distribution of log_*e*_(height) *y* at time *t* + 1 . The expression *P*(*y,x*; **θ**(*t*)) + *F*(*y,x*;**θ**(*t*)) is called the kernel, *K*(*y,x* **θ**(*t*)), which is a non-negative surface describing all possible demographic transitions from log_*e*_(height) *x* at time *t* to log_*e*_(height) *y* at time *t* + 1. The function *P*(*y, x*; **θ**(*t*)) comprises the survival-growth component of a time-varying IPM and can be decomposed into two functions that determine the probability of survival of an individual at log_*e*_(height) *x, s*(*x*; **θ**(*t*)), and the likelihood that the individual will grow from log_e_ (height) *x* to log_e_(heigh) *y* over a year, *g* (*y, x*; **θ**(*t*)), such that *P* (*y, x*; **θ**(*t*)) = *s*(*x*; **θ**(*t* )) *g* (*y, x*; **θ**(*t*)). The *F* (*y, x*; **θ**(*t*)) function comprises reproductive component of a time-varying IPM and can be decomposed into four functions that determine the probability of reproduction of an individual at log_e_ (height) *x, p*_*f*_ (*x*; **θ**(*t*)), the number of inflorescences/spikes of an individual at log_e_ (height) *x, f*_*n*_ (*x*; **θ**(*t*)), the probability a recruit establishes *p*_*e*_ (**θ**(*t*)), and the probability distribution of recruit size *f*_*d*_ (**θ**(*t*)), such that *F* (*y, x*; **θ**(*t*)) = *p*_*f*_ (*x*; **θ**(*t*)) *f*_*n*_ (*x*; **θ**(*t*)) *p*_*e*_ (*x*; **θ**(*t*)) *f*_*d*_ (**θ**(*t*)).

To assess the effects of grazing intensity on the long-term population persistence in fluctuating environments, we calculated the stochastic population growth rate (λ_S_) under each grazing intensity per species. The λ_S_ is given by the long-run geometric mean of annual growth rates (Rees & Ellner 2009)

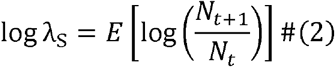

Here, *N*_*t*_ is the total population size summed across sizes at time *t* and *E* represents the expected value. The λ_S_ indicates whether a population will increase (λ_S_ > 1) or decline to extinction (λ_S_ < 1) for a long time (Tuljapurkar 1990). We first discretized the time-varying IPM kernel into four large transition matrices of size 200 × 200 using the midpoint rule (Ellner & Rees 2006). Using a sequence of Markovian environmental states derived from these matrices (Appendix S1: Figure S5), we then simulated population dynamics for 10,000 years. To exclude transient dynamics, we calculated the geometric mean growth rate of the latter *T* = 9,000 years, and computed 95% confidence intervals of λ_S_ (Caswell & Kaye 2001; Elderd & Miller 2016). To further evaluate the effect of grazing on *λ*_S_, we calculated the difference in logλ_S_ values (Δlogλ_S_) between grazing treatments and grazing exclusion.

To assess the effects of environmental variation on the population performance under each grazing intensity, we calculated the annual population growth rate (λ) for each environmental condition and quantified the temporal variance in annual population growth rate (τ^2^) among these environments. The λ informs on whether a population will increase (λ *>* 1) or decline to extinction (λ *<* 1) under a stationary environment. To estimate the uncertainty in λ, we bootstrapped the demographic data 1,000 times to obtain possible λ. The τ^2^ was calculated using Tuljapurkar’s small noise approximation (Tuljapurkar 1982; Rees & Ellner 2009)

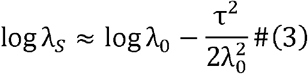

where, λ_0_ is the dominant eigenvalue of the mean kernel ***K***_*t*_, ***K***_*t*_ is shorthand for *K* (*y, x*; **θ**(*t*)). An increase in τ^2^ leads to reduction in λ_S_ (Hilde et al. 2020). The second term in Eq. (3) represents the contributions of temporal variation in vital rates and can be further decomposed into summed variance contributions of individual vital rates and summed covariance contributions among vital rates (H). Values of H showed a balance between that the contributions of negative and positive vital rate correlation (Jongejans et al. 2010; Gascoigne et al. 2025). To evaluate whether the demographic responses to grazing may be affected by species’ life-history traits, we calculated mean age at first reproduction, mean lifespan from reproduction, and overall lifespan based on the mean transition matrix for each species, using the “*Rage*” package (Jones et al. 2021) in R.

### 2.6 Stochastic elasticity to population growth

To examine the relative importance of each demographic process to the long-term population persistence, we conducted stochastic elasticity analyses of λ_S_ to each vital rate. This approach quantifies how λ_S_ responds proportionally to small changes in individual vital rates in a temporally variable environment (Caswell 2005). The elasticity of λ_S_ to specific vital rate *θ* is calculated

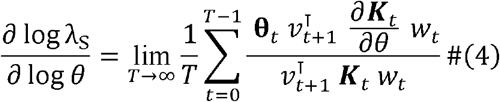

Here, *w*_*t*_ and *v*_*t*+1_ are the time-varying population structure and reproduction value, respectively, and **θ**_*t*_ is shorthand for **θ**(*t*). To assess the effects of both mean vital rates and demographic variability on stochastic population growth, We calculated the elasticities of λ_S_ to the mean vital rate (*e*^µ^) and to the standard deviation of vital rates (e^σ^) (Haridas & Tuljapurkar 2005; Rees & Ellner 2009).

### 2.7 Stochastic life table response experiment

To evaluate how vital rates contributed to grazing-induced changes in population growth, we performed stochastic life table response experiment (SLTRE) analyses. The SLTRE approximately decomposed the differences in stochastic population growth rate between grazing treatment *m* and grazing exclusion *r* (Δlogλ_S_^(m)^ into contributions from the differences in the means α and the temporal variability (*i*.*e*., standard deviations) β of vital rates (Davison et al. 2010):

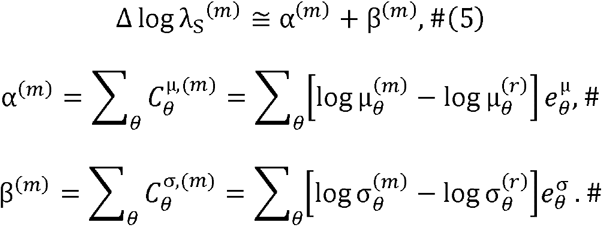

Here, 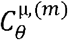 and 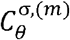 represent the contributions of differences in the means (μ) and the standard deviations (σ) of specific vital rates θ between grazing treatment and grazing exclusion. The 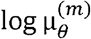 and 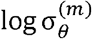 represent the natural logarithm of the means and the standard deviations of specific vital rate θ across years under grazing treatment, respectively, while 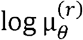 and 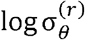 represent the corresponding values under grazing exclusion. The 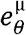 and 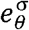are the elasticities to the means and to the standard deviations of specific vital rate θ for the halfway transition matrix between grazing treatment *m* and grazing exclusion *r* (Caswell 1996). To compare the relative vital rate contributions, we calculated the percentage contribution of each vital rate under each grazing treatment (*i*.*e*., 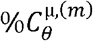 and 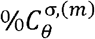), relative to the sum of absolute values of all contributions (Villellas et al. 2013). To evaluate the average effects of differences in each vital rate among grazing treatment, we calculated the percentage contributions of each vital rate across grazing treatment (*i*.*e*., 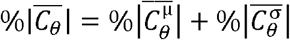). Finally, we compare the average effects of differences in means and standard deviations by calculating the percentage contributions of means 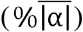 and standard deviations 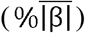 across grazing treatment (Appendix S2).

### 2.8 Test for demographic compensation and its effectiveness

To assess whether demographic compensation occurs in species’ response to grazing intensity, we used a randomization approach to evaluate whether the observed data harbored more negative correlations among sensitivity-weighted vital rates (*i*.*e*., vital rate contributions) than expected by chance (Villellas et al. 2015). We tested for demographic compensation by calculating negative Spearman’s correlations between pairs of vital rate contributions. The occurrence of a significantly higher number of significant negative correlations than expected by chance is indicative of demographic compensation and was tested by permuting the vital rate contributions among grazing intensities 1000 times (Andrello et al. 2020). To better understand the pattern of demographic compensation, we also examined how vital rate contributions changed with grazing intensity. We fitted linear regression models — with and without quadratic terms—relating each vital rate contribution to grazing intensity (measured as feeding rate, %), and selected the best-fitted models using the Akaike information criterion (Sheth & Angert 2018).

To further assess the effectiveness of demographic compensation in buffering the variation in population growth across grazing intensities 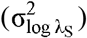, we conducted additional randomization tests. We recalculated 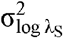 following the permutation of one vital rate contribution at a time (thus 6 vital rates). Higher values of 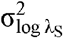 in the randomization indicated that the focal vital rate reduced 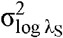 through its negative correlations; conversely, lower values for 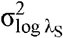 indicated the focal vital rate increased 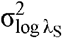 through its positive correlations (Andrello et al. 2020). Finally, we ran a targeted randomization test where we permuted only those vital rate contributions that reduced 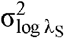: this randomization procedure eliminates as much as possible the negative correlations while preserving the important positive correlations and indicates the overall effectiveness of demographic compensation (Villellas et al. 2015).

## RESULTS

### Impacts of grazing on demographic processes

To examine how grazing affects demographic processes in alpine species with differing palatability, we constructed multiple mixed-effects models relating vital rates to grazing intensity and the natural logarithm of current plant height in *Morina chinensis* (less palatable) and *Deyeuxia flavens* (more palatable) (Table 1; Appendix S1: Table S1). We then performed model selection to identify the best-supported model for each vital rate in both species (Appendix S1: Table S2). The selected models revealed that grazing intensity had a strong influence on vital rates, with effects differing between species and varying across years (Appendix S1: Table S3).

The survival response to grazing differed between *M. chinensis* and *D. flavens* (Figure 2a, b). Light grazing consistently resulted in the greatest increase in survival across all plant sizes of *M. chinensis*, while in *D. flavens* it slightly increased the survival of taller individuals but decreased that of shorter ones (Figure 2a, b). Moderate grazing also increases in the survival of *M. chinensis* across all plant sizes, while in *D. flavens* it increased the survival of shorter individuals but slightly decreased that of taller ones (Figure 2a, b). Similarly, heavy grazing slightly increased the survival of *M. chinensis* across all plant sizes, yet in *D. flavens* it caused the greatest decline in survival of taller individuals and the greatest increase of shorter ones (Figure 2a, b).

**Figure 2.**
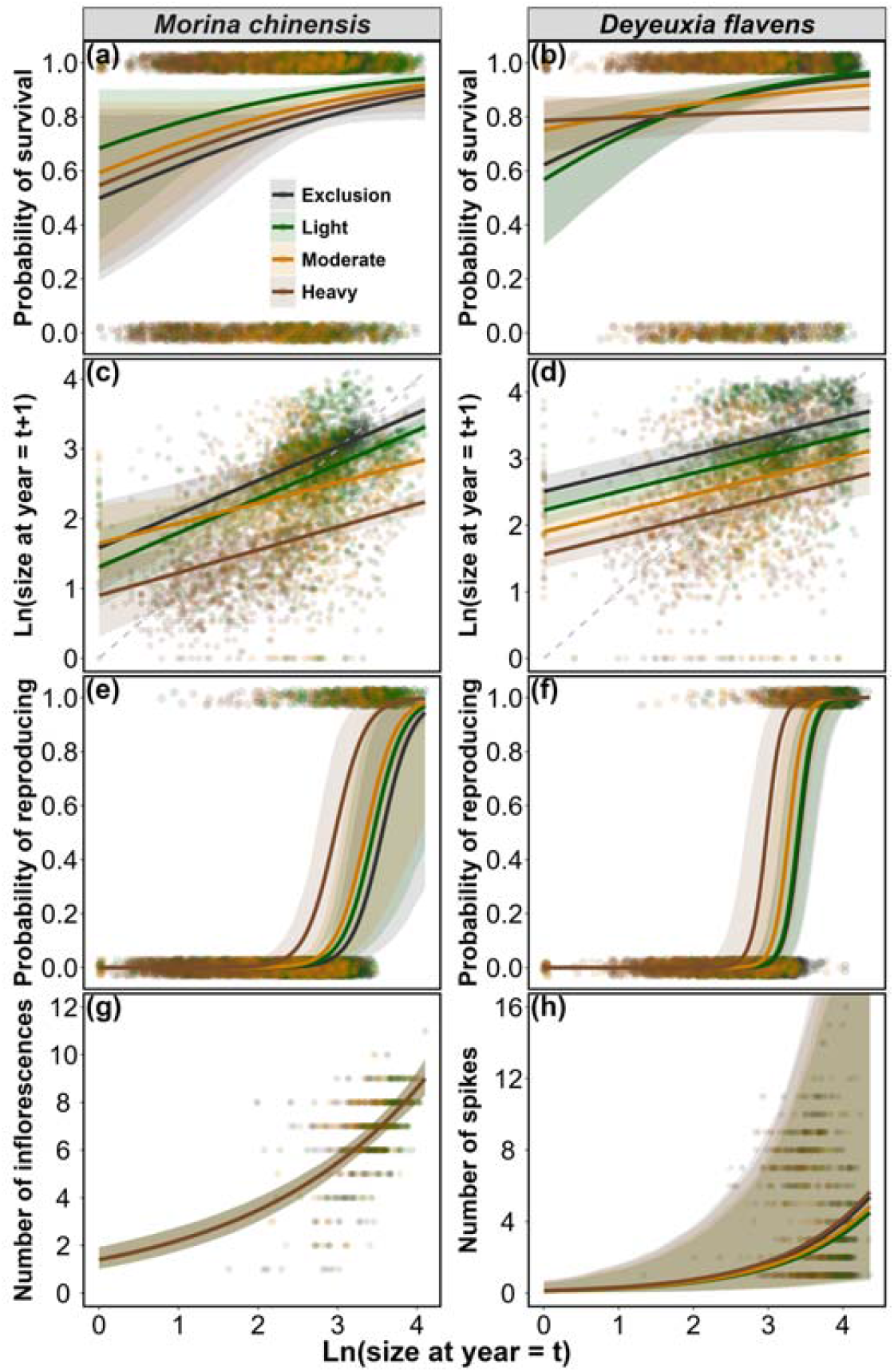
The effects of grazing intensity on the relationships between the vital rates of survival (a, b), growth (c, d), reproduction (e, f), and number of inflorescences/spikes per capital (g, h), and the natural logarithm of plant height during 2019-2023 for two co-occurring alpine herbaceous species, *Morina chinensis* (a, c, e, g) and *Deyeuxia flavens* (b, d, f, h), on the Tibetan Plateau. Colored solid lines represent the fixed effects of the natural logarithm of plant height on each vital rate across different grazing intensities, with 95% confidence intervals in the shaded areas (based on best-supported mixed-effects models; Appendix: Table S3). Colored points represent observed values of each vital rate at different grazing intensities. The grey dashed lines in (c) and (d) represent 1:1 line.

Grazing had a detrimental effect on changes in plant size of both species, with these negative effects intensifying progressively as grazing intensity increased (Figure 2c, d). Overall, changes in the natural logarithm of current plant height shifted from positive growth in shorter plants to shrinkage in taller plants, as indicated by the fitted regression lines rising above the 1:1 line for shorter plants and falling below it for taller plants (Figure 2c, d). Moreover, the degree of shift towards shrinkage intensified progressively with increasing grazing intensity in both species (Figure 2c, d).

Grazing consistently increased the probability of reproduction across all plant sizes for both species, with these effects intensifying as grazing intensity increased (Figure 2e, f). Specifically, the threshold of the natural logarithm of current plant height at which individuals reached a 50% probability of reproduction increased with increasing intensity in both species (Figure 2e, f). However, the response of seed production to grazing was different between species: the number of inflorescences remained unchanged across grazing intensities in *M. chinensis*, whereas in *D. flavens* light and moderate grazing reduced the number of spikes in taller plants, while heavy grazing led to an increase (Figure 2g, h). Furthermore, light grazing resulted in the greatest increase in the seedling recruitment for *M. chinensis*, while seedling recruitment in *D. flavens* increased progressively with grazing intensity (Appendix S1: Figure S6a, b). In contrast, recruitment size decreased with increasing grazing intensity in both species (Appendix S1: Figure S6c, d).

#### Impacts of grazing on population performance

To evaluate how grazing-induced changes in demographic processes under varying environmental conditions scale up to affect the overall population performance of both study species, we built stochastic integral projection models (SLTRE) to estimate the stochastic population growth rate (λ_S_), the temporal variance in annual population growth rate (τ^2^ ) under each grazing intensity, and the deterministic population growth rate (λ) under each environmental condition. Our SLTRE results revealed a tipping point in λ _S_ along the grazing intensity gradient for both species: light grazing consistently maintained the highest λ_S_, moderate grazing led to a reduction, and heavy grazing resulted in the lowest λ_S_ (Figure 3a, b). In parallel, τ^2^ increased with increasing grazing intensities for both species, with τ^2^ for *D. flavens* consistently lower than that for *M. chinensis* at each grazing intensity (Figure 3c, d). Furthermore, λ varied across environmental conditions, and these variations differed among grazing intensities for both species (Appendix S1: Figure S7a-h). Interestingly, although *D. flavens* had a later age at first reproduction than *M. chinensis*, it exhibited both a longer reproduction lifespan and a longer total lifespan (Table 1).

**Figure 3.**
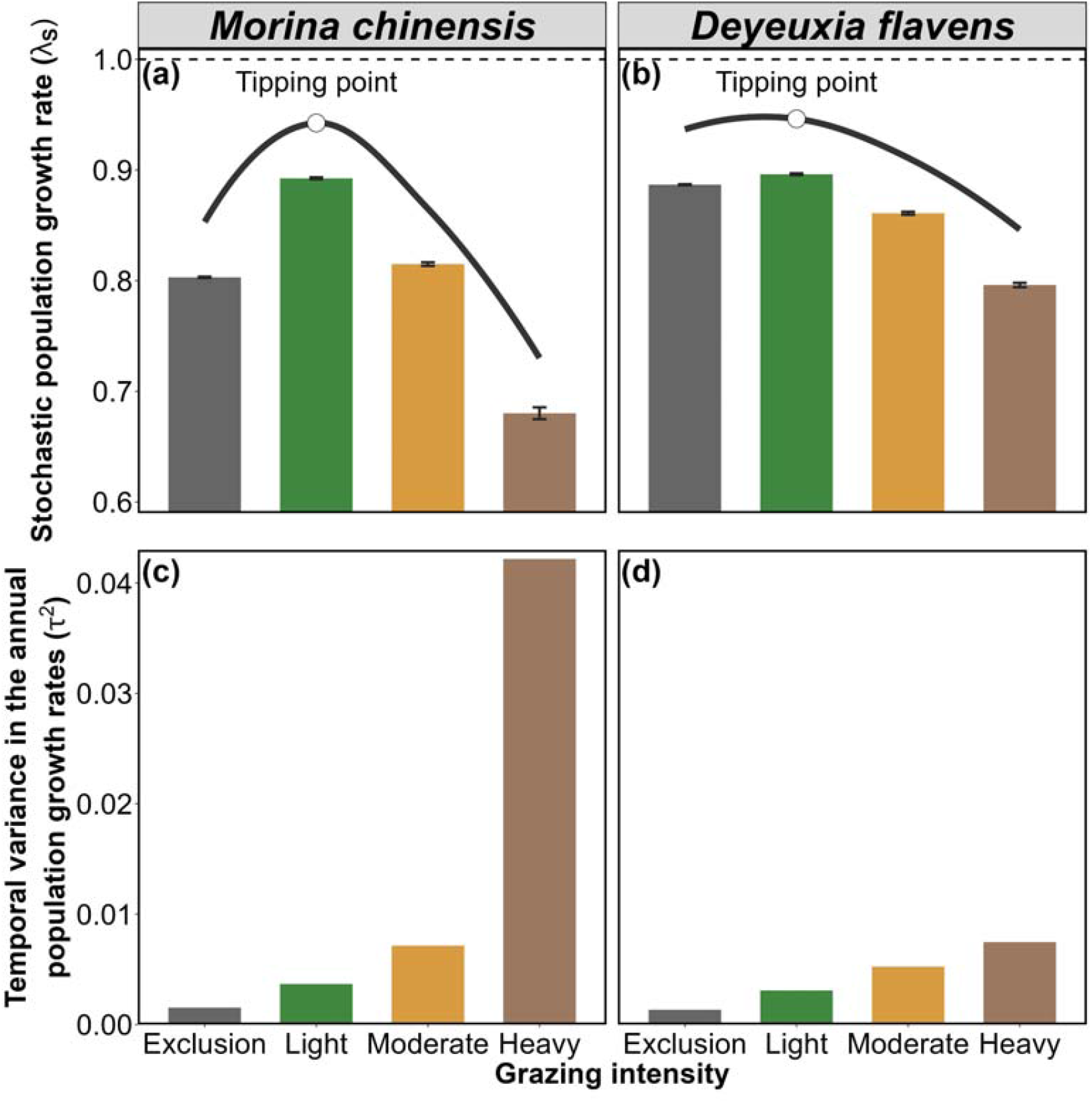
The effect of grazing intensity on the stochastic population growth rates (λ_S_) (a, b) and the effect of environmental stochasticity on the temporal variance in the annual population growth rate (τ^2^) during 2019-2023 under each grazing intensity (c, d) for two co-occurring alpine species *Morina chinensis* (a, c) and *Deyeuxia flavens* (b, d), on the Tibetan Plateau. The horizontal dashed line at 1 in (a, b) represents the baseline level for stochastic population increase (λ_S_ > 1) or decline to extinction (λ_S_ < 1). The error bars in (a, b) represent 95% confidence intervals of λ_S_.

#### Impacts of grazing on elasticity of demographic processes to population growth

To evaluate the relative importance of each demographic process to the population performance of both study species, we performed stochastic elasticity analyses of stochastic population growth rate to the mean (*e*^µ^) and standard deviation (*e*^n^) of each vital rate. Our results showed that λ_S_ was overwhelmingly elastic to changes in the mean survival rate across all grazing intensities in both species (Figure 4). Notably, the *e*^µ^ of survival increased towards shorter individuals but decreased towards taller individuals with increasing grazing intensity (Figure 4). The summed *e*^µ^ values of growth, fecundity, seedling recruitment and recruit size were consistently positive but considerably smaller than that of survival, whereas the summed *e*^µ^ values of shrinkage remained consistently negative (Figure 4). Furthermore, the magnitudes of *e*^n^ were generally much smaller than those of *e*^µ^ (Figure 4).

**Figure 4.**
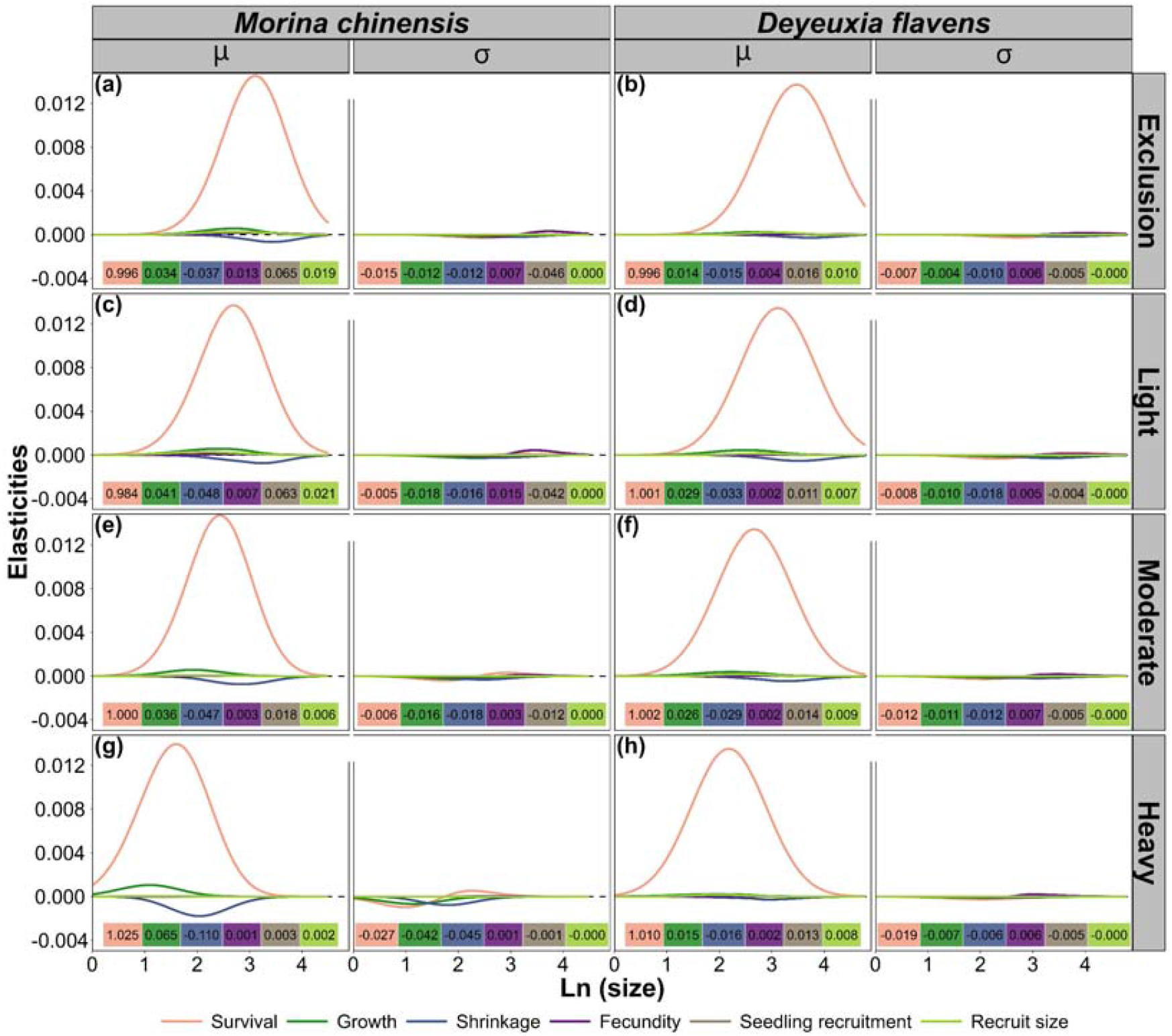
The effect of grazing intensity on the elasticities of stochastic population growth rate to changes in the mean (μ) and the standard deviation (σ) of each vital rate for two co-occurring alpine herbaceous species, *Morina chinensis* (a, c, e, g) and *Deyeuxia flavens* (b, d, f, h), on the Tibetan Plateau. Elasticity values for the mean and standard deviation of each vital rate accumulated through all sized are given in the tables at the bottom of each panel.

#### Demographic drivers of grazing-induced variation in population growth

To quantify the demographic processes underlying grazing-induced changes in population growth under varying environmental conditions for both study species, we performed a stochastic life table response experiment (SLTRE) analysis to decompose the differences in stochastic population growth rates (λ_S_) between grazing treatments and grazing exclusion (Δlogλ_S_) into contributions from the differences in the means (α) and the standard deviations (β) of each vital rate. Overall, differences in means had the greatest influence on Δlogλ_S_ for both species 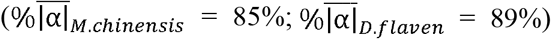, but the effects of standard deviations were not negligible 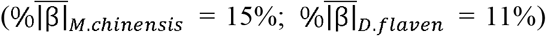 and could account for up to 50% of Δlogλ_S_ under specific grazing intensities (Figure 5b, c). Regarding contributions of vital rates to Δlogλ_S_, the largest were attributed to survival, growth and shrinkage, whereas reproductive vital rates had smaller contributions (Appendix S1: Figure S8).

**Figure 5.**
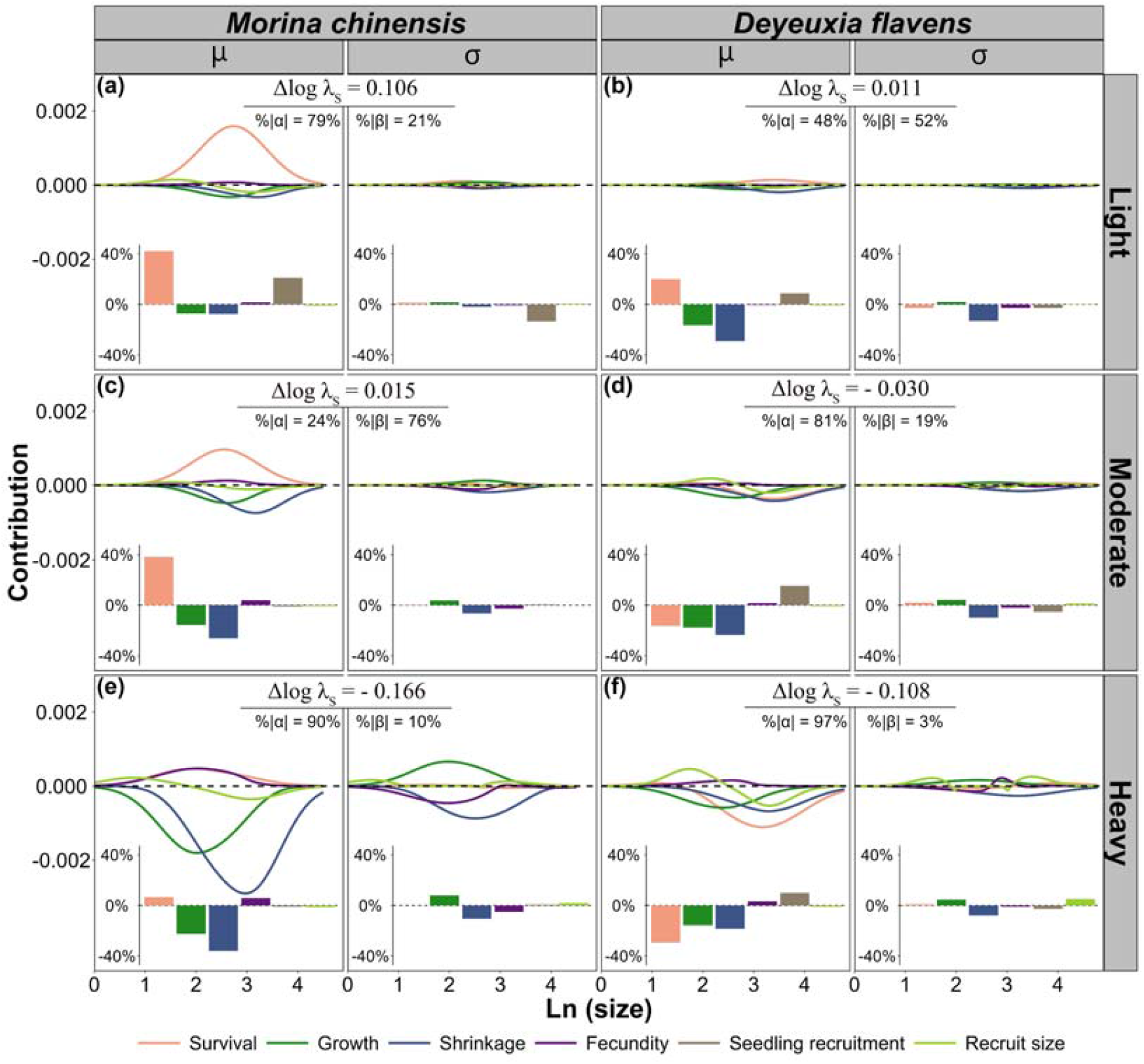
Results of stochastic life table response experiment (SLTRE) analyses to examine the effects of grazing intensity on the contributions of differences in mean values (μ) and of differences in the standard deviation (σ) to the changes in stochastic population growth rate between grazed and ungrazed populations (Δlogλ_S_) of two co-occurring alpine herbaceous species, *Morina chinensis* (a, c, e) and *Deyeuxia flavens* (b, d, f), on the Tibetan Plateau. The percentage net contributions of the means (%|α|) and the standard deviations (%|β|) of all the vital rates for grazed populations are given at the top of each panel. The percentage contribution of means and standard deviations of each vital rate for grazing populations, relative to the sum of absolute values of all contributions, were calculated and are given in the bar chart at the bottom of each panel.

The increase in λ_S_ under light grazing for both species was primarily driven by enhanced survival of taller individuals and increased seedling recruitment (Figure 5a, b). Under moderate grazing, the slight increase in λ_S_ for *M. chinensis* was mainly due to improved survival of taller individuals, whereas the slight decline in λ_S_ for *D. flavens* was primarily caused by increased shrinkage, suppressed growth, and reduced survival of taller individuals (Figure 5c, d). Under heavy grazing, the severe decline in λ_S_ for *M. chinensis* was largely attributed to greater shrinkage and suppressed growth of taller individuals, while the moderate decline for *D. flavens* was largely caused by reduced survival, intensified shrinkage, and decreased growth in taller individuals (Figure 5e, f).

#### Demographic compensation alleviates heavy grazing-induced population decline

To examine whether different demographic responses to grazing intensity can buffer associated variation in population growth 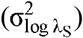, we used a randomization approach to test for demographic compensation and its effectiveness. In *D. flavens*, we found eight significantly negative correlations out of 15 possible pairwise correlations across grazing intensities (Figure 6b). This number was significantly greater than expected by chance (*P* = 0.02; Figure 6b), indicating the presence of demographic compensation in *D. flavens*. Fecundity and seedling recruitment contributed most to this demographic compensation (Figure 7a), although their effectiveness in reducing 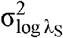 was relatively limited (Figure 7b). In contrast, the number of negative correlations across grazing intensities in *M. chinensis* was not significantly greater than expected by chance (*P* = 0.87; Figure 6a). Additionally, we found that summed contributions of the temporal covariance among vital rates were lower in *D. favens* across all grazing intensities (Appendix S1: Figure S9), indicating more negative covariances between vital rates in *D. favens* than in *M. chinensis*.

**Figure 6.**
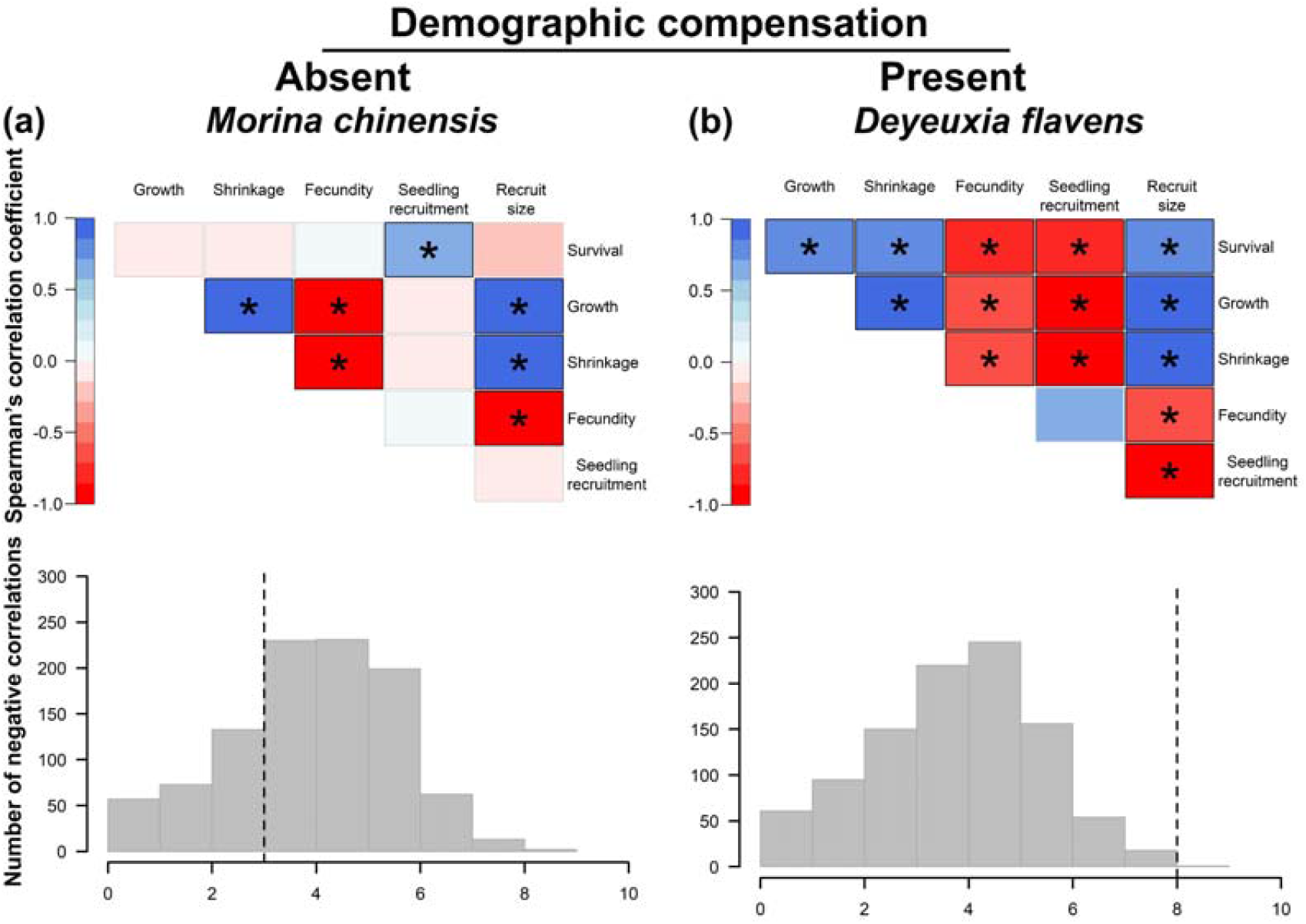
The test for demographic compensation among vital rates across the grazing intensity for the two co-occurring alpine herbaceous species, *Morina chinensis* (a) and *Deyeuxia flavens* (b), on the Tibetan Plateau. Correlogram of spearman’s correlation coefficients between the contributions of different vital rates to differences in stochastic population growth rates among the grazing intensity. Negative correlations are in red, positive correlations are in blue. Boxes with thicker borders and an asterisk indicate significant correlations at *P* < 0.05. The histograms show the number of observed significant negative correlations (vertical dotted lines) compared to the distributions of the number expected by change obtained through a permutation test.

**Figure 7.**
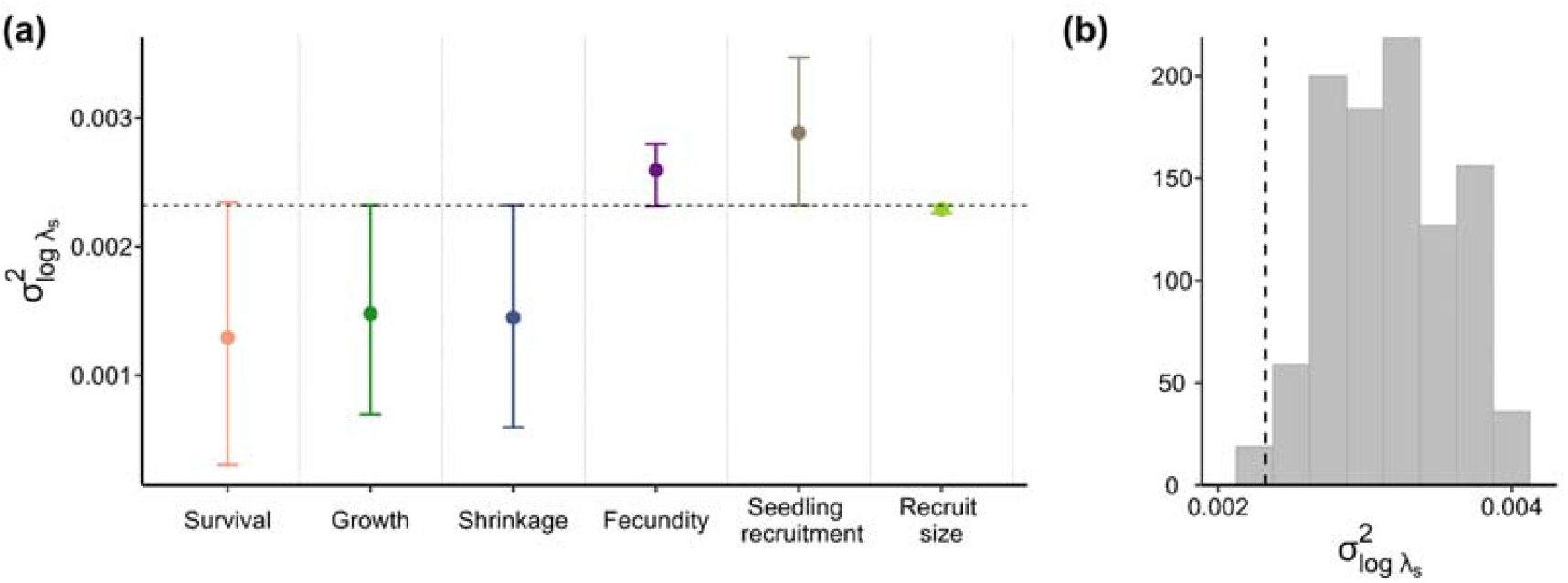
Effectiveness of demographic compensation in *Deyeuxia flavens*. (a) variance in population growth rate log λ_S_ among the grazing intensity 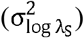 obtained by permuting each vital rate at a time. Dots are mean values and bars extend over 95% confidence intervals over 1000 permutations. Values higher than the observed 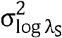 (horizontal dashed line) indicated that the corresponding vital rates effective reduces 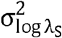 through demographic compensation. (b) observed 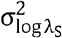 (vertical dashed line) compared to the distribution of 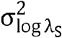 expected by chance obtained through permuting only the vital rate effectively reducing it (histograms).

To assess how demographic compensation buffers population decline under heavy grazing, we used linear regression to evaluate how vital rate contributions varied with grazing intensity. We found that the population-specific contributions of vital rates to population growth varied across grazing intensities and differed between species (Figure 8a, b; Appendix S1: Table S4). In *D. flavens*, the declines in contributions of survival, growth, and shrinkage towards higher grazing intensity were partially offset by increases in contributions from fecundity and seedling recruitment, thereby buffering the population decline under heavy grazing (Figure 8b). In contrast, the reductions in growth and shrinkage contributions towards higher grazing intensity were too substantial to be compensated solely by increased contributions from fecundity in *M. chinensis* (Figure 8a).

**Figure 8.**
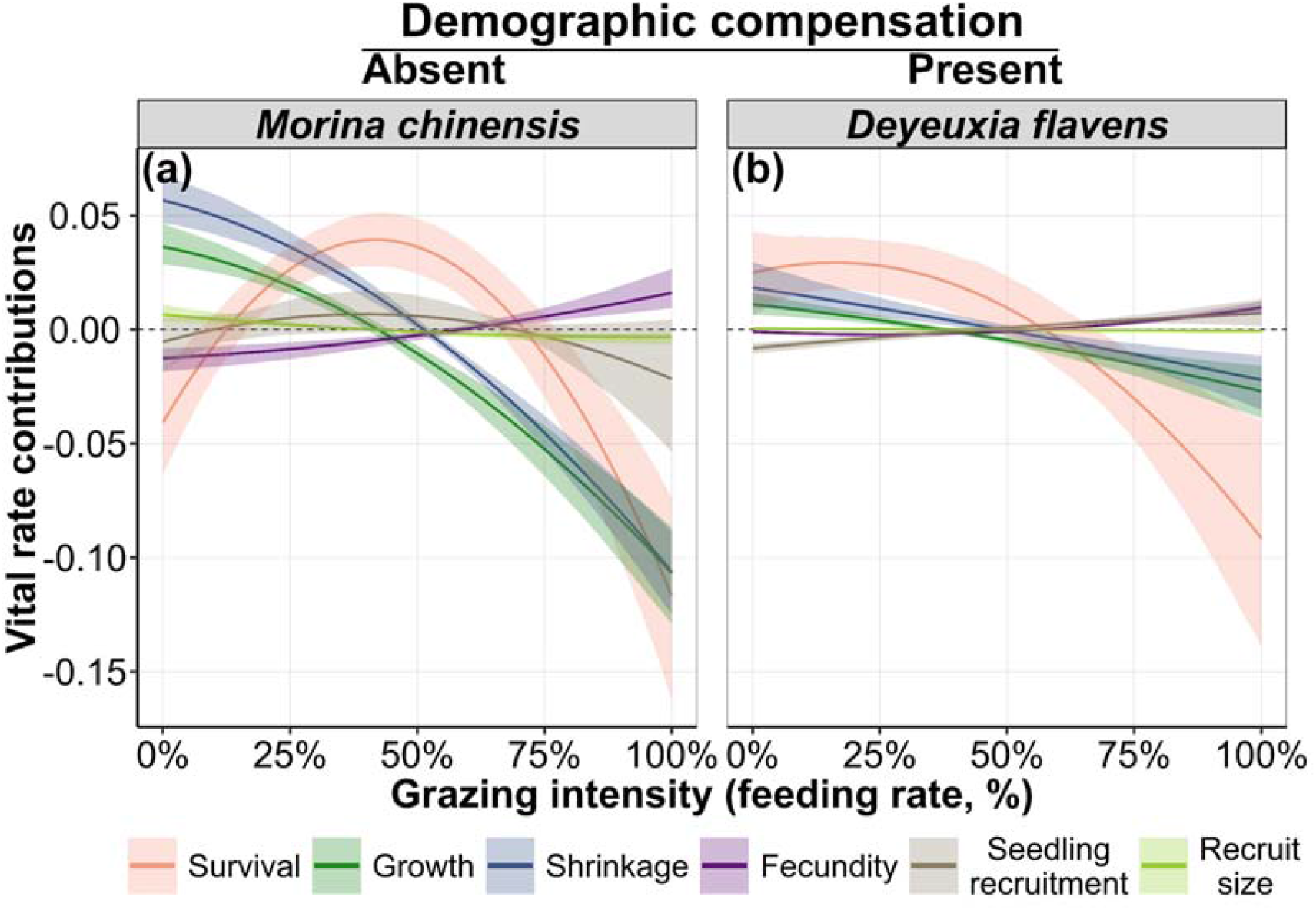
Population-specific vital rate contributions as functions of grazing intensity (feeding rate, %) of two co-occurring alpine herbaceous species, *Morina chinensis* (a) and *Deyeuxia flavens* (b), on the Tibetan Plateau. Colored solid lines represent linear and quadratic terms, with 95% confidence intervals in the shaded areas (Appendix S1: Table S5).

## DISCUSSION

Determining the optimal grazing intensities for the long-term population maintenance is crucial for informing management policy (Menges 2000; Bardgett et al. 2021; Larios & Hallett 2022). It requires long-term demographic studies integrating vital rates over an entire life cycle across well-controlled gradients of grazing intensity (Bastrenta et al. 1995; Larios & Hallett 2022). Here, we conducted a five-year consecutive demographic study and parameterized stochastic integral projection models (stochastic IPMs) to examine how different grazing intensities affect the vital rates across the entire life cycle and ultimately shape the long-term population dynamics for tow coexisting herbaceous species on the Tibetan Plateau grasslands. We uncover a hump-shaped relationship between the stochastic population growth rate and grazing intensities, providing the first population-level empirical evidence in support of the ‘Intermediate Disturbance Hypothesis’. Importantly, we demonstrate that demographic compensation among vital rates helps to mitigate population decline under heavy grazing. Furthermore, we show that temporal variance in vital rates—often overlooked—can rival the effects of mean vital rates, and markedly reduce population growth under heavy grazing pressure. Together, our findings provide mechanistic insights into the key demographic processes driving population dynamics under a gradient of grazing intensity, thereby better informing sustainable management strategies.

### Hump-shaped response of population growth to grazing intensities

Community studies have frequently reported the peak biodiversity at intermediate grazing intensity (Huston 1979; Miller et al. 2011; Hart & Jenkins 2023), supporting the ‘Intermediate Disturbance Hypothesis’ (Grime 1973; Connell 1978), but it remains unknown whether this hypothesis still holds at population level. We found that the stochastic population growth rate (λ_S_) of both study species initially increase with grazing intensity, but subsequently decline as grazing pressure intensifies, ultimately reaching the lowest value under heavy grazing. The enhanced λ_S_ under low grazing intensity was primarily attributed to the increase in both survival and seedling recruitment of both study species, while the reduced *λ*_S_ under high grazing intensity was caused by diminished growth and intensified shrinkage. Similar hump-shaped responses to grazing pressure have been widely reported across multiple scales, including traits (Liu et al. 2019; Shi et al. 2023), individual performance (Wang et al. 2016; Oñatibia & Aguiar 2019), and ecosystem functions (Ren et al. 2018; Wu et al. 2025). For example, Liu *et al*. (2019) found that the specific leaf area of *Leymus chinensis* increased at low grazing intensity and then declined toward high grazing intensity, probably because low grazing promotes rapid growth of thin leaves for efficient light capture (Poorter et al. 2010), whereas high grazing induces thicker, more durable leaves that resist herbivore damage (Villar et al. 2013); Wang *et al*. (2016) found that the aboveground biomass of *Kobresia humilis* initially increased at light grazing, likely because light grazing reduced shading by taller species that allowing higher leaf photosynthetic rates (Thomas & Bazzaz 1999), whereas at higher grazing intensities, biomass declined due to reductions in leaf area and leaf biomass, which limited photosynthesis and overall plant productivity (Milla & Reich 2007); Ren *et al*. (2018) found that ecosystem multifunctionality of semi-arid grassland was maximized at moderate grazing intensity, primarily because this grazing pressure maintained the highest aboveground net primary productivity (Liu et al. 2015; Li et al. 2017). These findings across different scales, together with our results, provide compelling evidence of the hump-shaped response of plants to grazing intensity.

### Demographic compensation buffers population decline under heavy grazing

Demographic compensation—the opposing responses of vital rates along grazing gradients—may buffer the detrimental effects of intense grazing on population maintenance (Villellas et al. 2015; Sheth & Angert 2018). In our study, although heavy grazing caused population declines in both species, the decline in *D. flavens* was markedly smaller than that in *M. chinensis*. Such contrasting population responses under high grazing pressure are largely attributable to whether demographic compensation occurs. Specifically, we found strong evidence of demographic compensation across grazing gradients in *D. flavens*, whereby reductions in survival, growth, and shrinkage toward higher grazing intensity were partially offset by increases in fecundity and seedling recruitment. These compensatory responses of vital rates alleviated population decline of *D. flavens* under high grazing pressure, whereas the absence of such compensation in *M. chinensis* resulted in a sharp decline in population growth. Demographic compensation buffering population decline towards high grazing pressure has also been documented across ecosystems (Evju et al. 2010, 2011; Johansen et al. 2016). For example, Johansen *et al*. (2016) showed that reductions in clonal reproduction and flowering probability under high grazing intensity was partially counterbalanced by enhanced sexual reproduction, thereby forestalling population decline of *Knautia arvensis* in the boreal vegetation zone. Similarly, Evju *et al*. (2011) found that the reduced survival with increasing grazing intensity was largely offset by increases in retrogression and clonal reproduction, resulting in stable population growth of *Geranium sylvaticum* in the low alpine zone. These findings, together with our results, suggest that demographic compensation can mitigate population decline under intense grazing pressure. Moreover, demographic compensation enabling population persistence under intense anthropogenic disturbance has been documented across gradients of clipping intensity (Lennartsson et al. 2018), leaf harvest intensity (Mandle et al. 2015), and fire severity (Hart-Fredeluces et al. 2021). As such, we add ourselves to the list of authors who have argued that species lacing demographic compensation across gradients of anthropogenic disturbance are more likely to face population decline under intense disturbance (Hegland et al. 2010; Xie et al. 2018).

Compensatory responses of vital rate to environment variation may buffer population decline under grazing (Jongejans et al. 2010; Gascoigne et al. 2025). Life-history theory predicts that negative covariances among vital rates reduce temporal variance in annual population growth rates, thereby stabilizing long-term population dynamics (Tuljapurkar 1982; Pfister 1998). Consistent with this prediction, regardless of grazing intensity, *D. flavens* exhibited more negative covariances among vital rates across years than *M. chinensis*, resulting in lower temporal variance in annual population growth rates. These compensatory responses of vital rates across years thus help buffer population decline of *D. flavens* under heavy grazing. Together, these theoretical predictions and our empirical results suggest that compensatory dynamics among vital rates help buffer populations against environmental fluctuations. Furthermore, our SLTER results revealed that changes in the temporal variance of vital rates contributed as much as shifts in their mean values to grazing-induced differences in population growth in both species. Similarly, Jacquemyn *et al*. (2010) found that temporal variability in vital rates had a non-negligible effect on grazing-induced changes in population growth of a perennial plant *Primula veris*. Collectively, these findings underscore the importance of incorporating temporal variability in vital rates when assessing population dynamics under anthropogenic pressures.

### Implications in grazing management

Livestock grazing is a common land-use activity in alpine grassland ecosystem (Bardgett et al. 2021; Zhang et al. 2023). We found population-level responses to grazing consistent with the ‘Intermediate Disturbance Hypothesis’ (Hickman & Hartnett 2002; Wang et al. 2016; Oñatibia & Aguiar 2019), suggesting grazing intensity must be managed within appropriate limits to maintain viable populations. Identifying which vital rates drive population maintenance is critical for effective grazing management (Haridas & Tuljapurkar 2005; Li et al. 2013). We found that survival was key for population maintenance of both species, suggesting that grazing should be controlled at levels that minimize mortality (Miao et al. 2025). Additionally, reduced growth made a considerable negative contribution to the population growth rates of both species in the current study, consistent with grazing-induced reductions in vegetative height and aboveground biomass on the Tibetan Plateau (Yang et al. 2016; Ji et al. 2020; Zhang et al. 2023). Therefore, these findings together with our results suggest that grazing activities should also be managed to a certain extent to avoid large decreases in growth. Given that survival and growth are relatively straightforward to monitor, they may be served as the key vital rates for the management of grazed populations in alpine grasslands.

Importantly, effective grazing management requires identifying species most vulnerable to grazing for conservation prioritization. We found that the population of *D. flavens*, which exhibits demographic compensation, declined less than *M. chinensis*, which lacks this trait, implying that conservation efforts should prioritize species without demographic compensation. Additionally, *D. flavens* exhibited lower temporal variance in vital rates and population growth than *M. chinensis*, thereby buffering its population against environmental fluctuations (Tuljapurkar 1982; Pfister 1998), suggesting that species with greater temporal variance in vital rates should be prioritized in conservation planning. Overall, species with demographic compensation and lower temporal variance are more resilient to environmental fluctuations and grazing, making them suitable for intensive grazing and restoration efforts, while species lacking these traits require targeted conservation to maintain viable populations.

## Author contributions

Shou-Li and Jin-Sheng He conceived the study. Hai-Tao Miao and Xiaofang Wang collected the data, Hai-Tao Miao constructed models and wrote first draft of the manuscript, and all authors contributed substantially to revisions. All authors approved the final manuscript.

## Findings

This work was funded by the National Natural Science Foundation of China (31971423) and the cost-share exchange projects between the National Natural Science Foundation of China and the Royal Society of the United Kingdom (32011530169).

## Data availability statement

The data and code that support the findings of this study are openly available in Zenodo at https://zenodo.org/records/17686600?token=eyJhbGciOiJIUzUxMiJ9.eyJpZCI6ImQ1_ZDhhYmEwLTI5NjAtNDlhYy05MGZmLWQzYTk1ZmViY2FlZCIsImRhdGEiOnt9_LCJyYW5kb20iOiI4MDU1OGU3YjJiN2FhOGZmYTZhMmViNzBjMDk3YTMxNi_J9.U5-vNTBvw9CYS_QXV83_T5w0F4Vk--jbxdPNxaZVnhi9io_aZCq0B6tfh3M-yuGjNoPTVxcENMxUcTSgSGEjXA.

